# Helminth infection modulates number and function of adipose tissue Tregs in high fat diet-induced obesity

**DOI:** 10.1101/2021.12.20.473610

**Authors:** Camila P. Queiroz-Glauss, Mariana S. Vieira, Marcela Helena Gonçalves-Pereira, Stephanie S. Almeida, Rachel H. Freire, Maria Aparecida Gomes, Jacqueline Isaura Alvarez-Leite, Helton C. Santiago

## Abstract

**Background:** Epidemiological and experimental studies have shown a protective effect of helminth infections in weight gain and against the development of metabolic dysfunctions in the host. However, the mechanisms induced by the parasite that regulate the development of metabolic diseases in the host are unclear. The present study aimed to verify the influence of *Heligmosomoides polygyrus* infection in early stages of high fat diet-induced obesity.

**Principal Findings:** The presence of infection was able to prevent exacerbated weight gain in mice fed with high fat diet when compared to non-infected controls. In addition, infected animals displayed improved insulin sensitivity and decreased fat accumulation in the liver. Obesity-associated inflammation was reduced in the presence of infection, demonstrated by higher levels of IL10 and adiponectin, increased infiltration of Th2 and eosinophils in adipose tissue of infected animals. Of note, the parasite infection was associated with increased Treg frequency in adipose tissue which showed higher expression of cell surface markers of function and activation, like LAP and CD134. The infection could also revert the loss of function in Tregs associated with high fat diet.

**Conclusion:** These data suggest that *H. polygyrus* infection can prevent weight gain and metabolic syndrome in animals fed with high fat diet associated with modulations of adipose tissue Treg cells.

**Author summary:** Helminth infections are known to modulate the immune system being responsible for protecting the host from developing allergic and autoimmune disorders (Hygiene Hypothesis). We hypothesized that the same immunomodulatory effect can have an impact on immunometabolic diseases, such as obesity and its linked diseases such as type 2 diabetes. Weight disorders have reached epidemic levels, nearly tripling since 1975 and being responsible for almost 5 million premature deaths each year. To test our hypothesis C57BL/6 male mice were fed control or high fat diet, for five weeks, in the presence or not of *Heligmosomoides polygyrus* infection. Weight gain, development of metabolic disorders, inflammation and cellular migration to the adipose tissue were evaluated. In accordance with our hypothesis, we found that the presence of infection prevented the exacerbated weight gain and also improved metabolic parameters in animals fed a high fat diet. This was associated with the infection’s ability to modulate parameters of a cell responsible for regulatory functions: Tregs. In the light of these findings, helminth infection could be protective against weight gain and metabolic disturbances.

## Introduction

Hygiene Hypothesis postulates that the stimulation of the immune system by microbes or microbial products, especially during childhood, can protect the host against the development of atopic and inflammatory disorders [1]. A number of epidemiological and experimental studies show a benefit effect of infections by viruses, bacteria, and helminths in the development of different inflammatory diseases like asthma [2 – 4], type 1 diabetes [5 – 8] and multiple sclerosis [9 – 10]. The positive influence of helminth infections in immunometabolic disorders, as obesity, has also been investigated [11 – 12].

Overweight and obesity are global health problems that reached epidemical status with more than 2 billion people with overweight and 650 million obese [13]. The greatest concern about this overnutrition syndrome is the association with the development of a number of serious metabolic consequences as glucose metabolism dysfunctions [12], heart diseases [14] and even some types of cancer [15 – 22]. The establishment of these dysfunctions along with weight gain is linked to the development of a low-grade chronic inflammation [12]. For example, adipose tissue fat accumulation is associated with the migration of inflammatory cells like macrophages [23 – 24], mastocytes [25] and T cells [26] that produce pro-inflammatory cytokines, such as TNF and IL6, which contributes to dysfunctions in glucose metabolism [27]. Interestingly, obesity has been associated with decreased frequency and dysfunction of adipose Treg cells, which is a possible mechanism to perpetuate background inflammation and metabolic syndrome [12, 28]. On the other hand, helminths are known to improve Treg function as a mechanism of immune evasion [29 – 31] and also in inflammatory disease models, such type 1 diabetes [25, 32] and asthma [33 – 34].

Helminth infections have been shown to influence the weight gain and the development of metabolic dysfunctions [12] in experimental models of obesity [35 – 41] and in epidemiological studies in humans [42 – 45]. Although these effects have been associated with increased eosinophils and Th2 cells infiltration and decreased Th1 and Th17 profiles [44, 46] in adipose tissue, the role of Tregs in helminth-obesity interface has been poorly investigated. The evaluation of Tregs is essential in obesity studies since their induction or transfer to obese animals is shown to inhibit metabolic syndrome parameters and it is discussed to be used as therapeutic strategy [28, 47, 48].

In the current study, we examined the effect of infection with *H. polygyrus* on early stages of high fat diet (HFD)-induced obesity in mice and showed the ability of the infection to modulate the number, phenotype and function of adipose tissue Tregs cells, which may play a role in preventing weight gain and metabolic dysfunctions. Our results show that improvement of metabolic syndrome associated with obesity experimental model is paralleled to improvement in Treg numbers and function in adipose tissue.

## Methods

### Animals, diets and *H. polygyrus* infection

Male C57BL/6Unib mice, specific pathogen free, with four weeks of age were obtained at the Central Breeding Center of the Federal University of Minas Gerais. Throughout the experiment, animals were housed in temperature-controlled room with 12-hour light-dark cycle, with food and water available ad libitum. Mice were fed with high fat diet (HFD) (62% energy derived from fat, 23% from carbohydrates and 15% from proteins) or low fat diet (LFD) (10% energy derived from fat, 74% from carbohydrates and 16% from proteins) for five weeks. Caloric intake was measured every week, considering the weight difference between the amount of diet offered and the next week’s left overs, divided by the number of animals per cage. This value was then multiplied by the number of calories per gram of each diet (LFD: 2.76 kcal/g; HFD: 5.21 kcal/g). Concomitantly with the diet, experimental group received 200 L3 *H. polygyrus* larvae by oral gavage. Eggs in feces were detected by visual observation of feces smears under microscope to confirm successful patent infection. Experiments were approved by the Ethics Committee for Animal Use of the Federal University of Minas Gerais (protocol#25/2012).

### Insulin tolerance test and oral glucose tolerance test

The insulin tolerance test (ITT) was performed four days before euthanasia. Animals were bled at the tail vein and glucose levels were measured by a glucometer (Accu-Chek Performa; Roche, Diagnostics, USA) before, and 15, 30 and 60 minutes after 0.75 U/kg insulin injection i.p. Two days before euthanasia, oral glucose tolerance test (OGTT) was carried out after 6 hours of fasting. Mice were given glucose (2g/kg of body weight) by gavage and blood glucose levels were measured by a glucometer (Accu-Chek Performa; Roche) before, and 15, 30, 60 and 120 minutes after gavage.

### Nutrition parameters and lipid profile analysis

At the end of five weeks, serum levels of total proteins, albumin, total cholesterol, HDL cholesterol and triglycerides were evaluated using commercial kits (Labtest Diagnóstica S.A., Brazil) after 16 hours fasting. Liver lipid quantification was analyzed according to the Folch method [49]. Briefly, frozen liver tissue (100mg) was homogenized in 950µL of chloroform:methanol (2:1). 200µL of methanol were added to the mixture and the samples were centrifuged. The supernatant was transferred to a weighted clean tube, then mixed with 400µL of saturated saline solution. After centrifugation of the mixture, the upper phase was discarded and the lower chloroform phase containing the lipids was washed three times with a Folch solution (2% NaCl 0.2%, 3% Chloroform, 47% distillated water, 48% methanol), evaporated under 60°C temperature, and total lipid weight was determined.

### Isolation of the Adipose Tissue Stromal Vascular Fraction

For isolation of the stromal vascular fraction (SVF) from adipose tissue [50], epididymal white adipose tissue (EWAT) was collected and then minced in DMEM, containing 4% Bovine Albumin Serum Fatty Acid Free and 0.1% glucose. Collagenase VIII (Sigma-Aldrich, Merck KGaA, USA) was added, at 4 mg/g of tissue, to the mixture containing the minced tissue followed by incubation at 37°C under constant agitation for 40 minutes. Following centrifugation, floating adipocytes were separated and the SVF pellet ressuspended and analyzed.

### Cell culture

Adipocytes (3×10^5^ cells/mL) and cells from SVF (5×10^6^ cells/mL) were cultured in 5% CO_2_ incubator at 37°C for 24 hours in the absence or presence of PMA (0.4mg/mL) and Ionomycin (5mg/mL). After incubation, the supernatant was collected and stored at −70°C until use. Cytokine levels were assessed by Cytometric Bead Array (CBA) Mouse Th1/Th2/Th17 (BD Biosciences, USA) and adiponectin secreted by the culture of adipocytes was assessed by ELISA (R&D System, USA).

### Flow cytometry

For analysis of eosinophils, SVF cells were stained with antibodies against CD11b (M1/70; BioLegend, USA), and Siglec F (E50-2440; BD Biosciences, USA) flowed by fixation with paraformaldehyde. Data from cell acquisition were analyzed and after defining the intersection population from the gates of leucocytes, single cells and time, cells stained with CD11b^int^SiglecF^+^ were considered eosinophils (S1 Fig).

For analysis of lymphocyte subsets, SVF cells were stained with antibodies against CD3 (17A2; BioLegend) and CD4 (GK1.5; BioLegend), then fixed and permeabilized with fix/perm buffer (eBioscience, Thermo Fisher Scientific, USA) according to the manufacture’s instructions. Cells were then incubated with antibodies against Tbet (4B10; BioLegend), RORgT (Q31-378; BD Biosciences) and Gata3 (16E10A23; BioLegend). Lymphocytes were first identified by forward/side scatter dot plot, then doublets and possible interruptions in the acquisition were excluded. Positive cells for CD3 and CD4 were selected. Then CD3^+^CD4^+^Tbet^+^ were considered Th1 cells, CD3^+^CD4^+^Gata3^+^ Th2 and CD3^+^CD4^+^RORgT^+^ Th17 (S2 Fig).

For characterization of Treg subtypes, SVF cells were stained with antibodies against CD3 (17A2; BioLegend), CD4 (GK1.5; BioLegend), GITR (YGITR 765; BioLegend), LAP (TW7-16B4; BioLegend), CD25 (PC61; BioLegend), CD134 (OX-86; BioLegend) and CD152 (UC10-4B9; BioLegend), then fixed and permeabilized with fix/perm buffer. Cells were then incubated with antibody against Foxp3 (150D; BioLegend). CD3^+^CD4^+^ T lymphocytes were selected as described above and then population positive for CD25 and Foxp3 was selected. CD3^+^CD4^+^CD25^+^Foxp3^+^ cells were considered Tregs and analyzed for surface marker (S3 Fig).

Flow cytometry data were collected on BD LSRFortessa™ Flow Cytometer using BD FACSDiva™ Software and gates were set according to unstained cells using FlowJo (version 10.5.3, Tree Star Inc, USA).

### In vitro Treg suppression assays

Tregs from EWAT were isolated from SVF cells using Dynabeads™ FlowComp™ Mouse CD4^+^CD25^+^ Treg Cells Kit (Invitrogen, Thermo Fisher Scientific, USA). T effector cells (Teff), isolated from the spleen of a control animal (non-infected and fed with regular chow) using the same kit, were stained with 5µM CFSE (Invitrogen, USA) for 10 minutes. Functional Treg assay was performed as described [51]. Briefly, Teff cells were co-cultured with Tregs at indicated proportions and stimulated with Dynabeads Mouse T-Activator CD3/CD28 (Life Technologies, USA) for 72 hours in 5% CO_2_ incubator at 37°C. Cells were acquired in BD LSRFortessa™ Flow Cytometer and the decay in CFSE fluorescence was analyzed with FlowJo (version 10.5.3, Tree Star Inc) using the tool Proliferation Modeling.

### Statistical analysis

All results were expressed as mean ± standard error of the mean. Group means were compared by Mixed-effects analysis or two-tailed Student’s test using GraphPad Prism (version 9.0.0, GraphPad Software Inc, USA). Probability values below 0.05 were considered statistically significant.

## Results

### *H. polygyrus* infection attenuated weight gain and metabolic dysfunctions

To determine the effect of *H. polygyrus* infection on early stages of obesity induced by high fat diet (HFD), C57BL/6 male mice were infected or not with L3 *H. polygyrus* larvae and fed with HFD diet for 5 weeks. The infection was able to prevent exacerbated weight gain in animals fed with HFD (Fig 1A) without decreasing the amount of food intake (Fig 1B). The lower weight gain in the HFD Hp group was associated with decreased weights of epididymal (EWAT) and subcutaneous (SAT) adipose tissue (Fig 1C). Importantly, this disparity in body weight was not observed between the groups that received control diet (Fig 1A), which suggests that the differences between infected and non-infected groups treated with HFD are not due to parasite spoliation of the host. Another data that suggests the host is not being spoliated by the parasite is that neither total serum protein or serum albumin levels differed between infected and non-infected animals (Figs 1D and E). Therefore, we conclude that *H. polygyrus* infection improved weight control in animals under HFD without causing malnutrition. Obesity is associated with dysregulation of glucose metabolism and hepatic steatosis (12) (27). Animals fed with HFD and infected with *H. polygyrus* presented lower fasting glycemic levels when compared to non-infected controls (Fig 1F). Further, HFD Hp group had a better response to glucose tolerance test showed by the faster return to blood basal levels after glucose injection, when compared to non-infected controls (Figs 1G and H). Both results indicate that the presence of the infection improved the development of glucose metabolic dysfunction associated to weight gain. Insulin sensitivity was also improved in infected animals when compared to the non-infected group. We found that blood glucose levels after insulin injection was higher in HFD Ni mice when compared to HFD Hp group, which suggested that the infection prevented the development of peripheral insulin resistance (Figs 1I and J). In addition, infected animals under HFD had significantly lower liver mass when compared to HFD Ni group (Fig 1K). This decrease in liver mass was directly associated with the amount of lipid/gram of liver, which was also lower in infected animals (Fig 1L). Furthermore, the infection also improved dyslipidemia associated with the HFD, i.e., decreased serum levels of triglycerides (Fig 1M) and also increased levels of HDL cholesterol (Fig 1N), besides not changing total cholesterol (Fig 1O). Overall, the effects of *H. polygyrus* infection on experimental obesity is in agreement with other models described in literature that helminth infection improves weight gain, fat accumulation and metabolic syndrome in animals fed with HFD [35 – 41].

**Fig 1.**
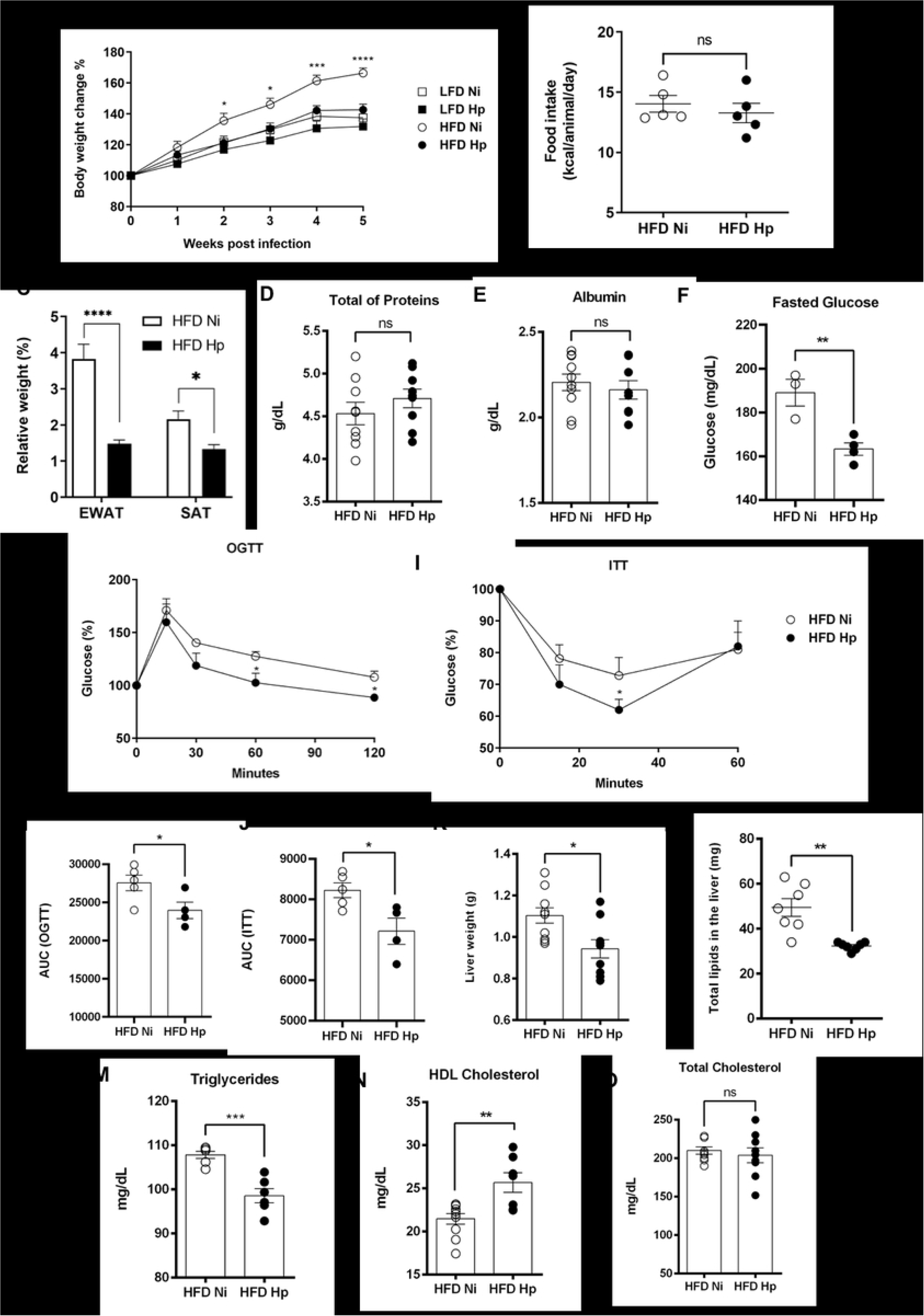
*H. polygyrus* infection attenuated weight gain and prevented metabolic dysfunctions in early stages of HFD-induced obesity. Animals were treated with control low fat diet (LFD) or with obesogenic high fat diet (HFD) and infected with *H. polygyrus* (Hp) or left uninfected (Ni). Body weight (A), caloric intake (B), relative weigh of epididymal white adipose tissue (EWAT), subcutaneous adipose tissue (SAT) (C) were assessed. Serum levels of total proteins (D) and albumin (E) were evaluated at the end of experiment. Fasting glycemic levels (F), response to oral glucose tolerance tests after glucose gavage (OGTT) (G and H) and insulin tolerance tests (ITT) after intraperitoneal injection of insulin (I and J) were compared between HFD-fed groups. Liver mass (K), hepatic quantification of lipids (L), fasting levels of serum triglycerides (M), HDL cholesterol (N) and total cholesterol (O) were assessed. N=4-10 representative of two or more experiments performed. Two-tailed T-Test was used to compare HDF Hp and HDF Ni groups. * p<0.05 ** p<0.01 *** p<0.001 **** p<0.0001 between HFD groups.

### *H. polygyrus* infection prevented the establishment of inflammation caused by high fat diet

The adipose tissue is an important endocrine organ that secrets a variety of hormones [52 – 53]. A key adipose tissue hormone is adiponectin, which is responsible for improving insulin function and plays many important homeostatic functions [54 – 55]. In addition, adipose tissue also secretes cytokines that are associated with the development of metabolic disorders [12]. Due to the importance of these adipokines and cytokines in the metabolism, we analyzed their production by EWAT. Adipocytes or SVF cells were separated and cultured for 24 hours, and adiponectin or cytokines were measured in the supernatants. We found no difference in the secretion of inflammatory cytokines like IL17A (Fig 2A), TNF (Fig 2B) and IL6 (Fig 2C) by SVF cells when comparing HFD Hp and HFD Ni groups. Production of IL2, IL4 and IFNγ were not detected in the culture of SVF cells (data not shown). On the other hand, the production of homeostatic adipocytokines like IL10, by SVF cells (Fig 2D), and adiponectin, measured in adipocytes supernatants (Fig 2E), was increased in HFD Hp group when compared to HFD Ni. These data suggest that the effect of *H. polygyrus* infection in metabolic parameters might be related to the secretion of anti-inflammatory or homeostatic adipocytokines, rather than a reduction of pro-inflammatory cytokines.

**Fig 2.**
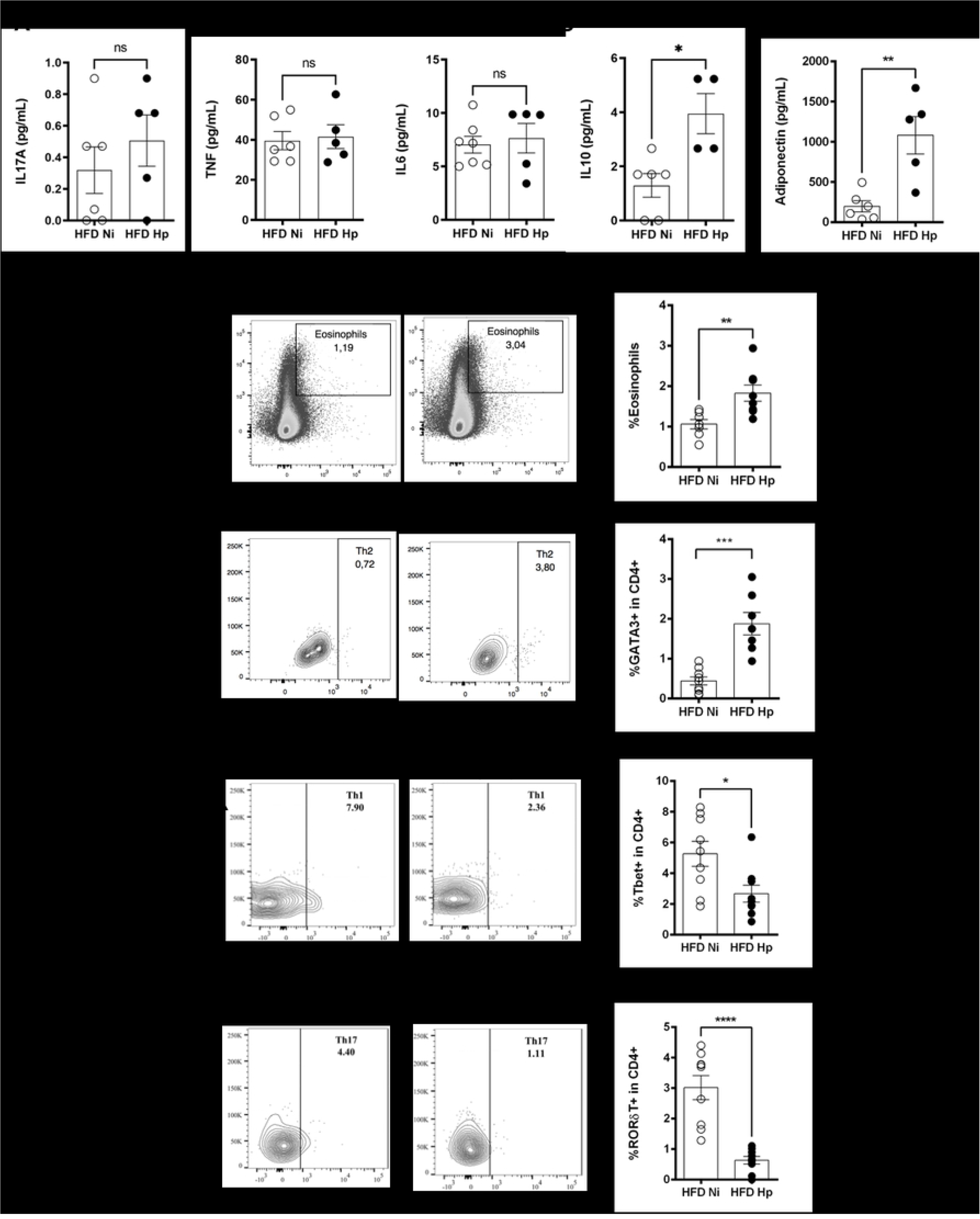
*H. polygyrus* infection prevented the establishment of inflammation caused by the high fat diet. Production of IL17A (A), TNF (B), IL6 (C) and IL10 (D) by stromal vascular fraction cells from epididymal white adipose tissue (EWAT). Production of Adiponectin by cultured adipocytes (E). Representative gates and frequency of eosinophils (F), Th2 (G), Th1 (H) and Th17 (I) cells isolated from EWAT and analyzed by flow cytometry. n=7-10 representative of two or more experiments performed. Two-tailed t test. * p<0.05 ** p<0.01 *** p<0.001 **** p<0.0001 between HFD groups.

To gain further insights about how the infection was modulating the inflammatory profile in adipose tissue, we evaluated the phenotype of the resident immune cells. We observed that the infection by *H. polygyrus* was associated with increased percentage of eosinophils (Fig 2F) and Th2 cells (Fig 2G) in the epididymal white adipose tissue. On the other hand, Th1 (Fig 2H) and Th17 (Fig 2I) cells were downmodulated by infection. Together, our data showed that the presence of the Hp infection is associated with an increased type 2 cells and decreased type 1 and type 17 infiltration in adipose tissue, despite maintenance of IL17A, TNF and IL6, that may have innate immune cells sources.

### *H. polygyrus* infection improved the number, phenotype and function of Tregs in adipose tissue

Obesity is associated not only with increased pro-inflammatory background, but also with Treg dysfunction [56 – 57]. On the other hand, despite the fact that helminth infections have been associated with improved Treg activity [25, 29, 33, 34, 58], their phenotype in an obesity-helminth infection interface has been poorly investigated. To gain insights about how helminth infection can influence the biology of Tregs in obesity, we analyzed the abundance of Tregs in adipose tissue and also the expression of cell surface markers in Tregs [28]. *H. polygyrus* infection increased the frequency of Tregs in adipose tissue (Fig 3A) in HFD-treated animals and modulated their phenotype. For example, the expression of LAP, a marker of membrane-bound TGF-ß [59], was increased in Tregs of Hp-infected animals when compared to HFD Ni group at MFI (Fig 3B) and frequency levels (Fig 3C). The infection also increased the frequency and per cell expression of CD134 by adipose tissue Tregs (Figs 3D and E). On the other hand, the expression of GITR and CTLA-4 by adipose tissue Tregs were not altered by the infection (data not shown).

**Fig 3.**
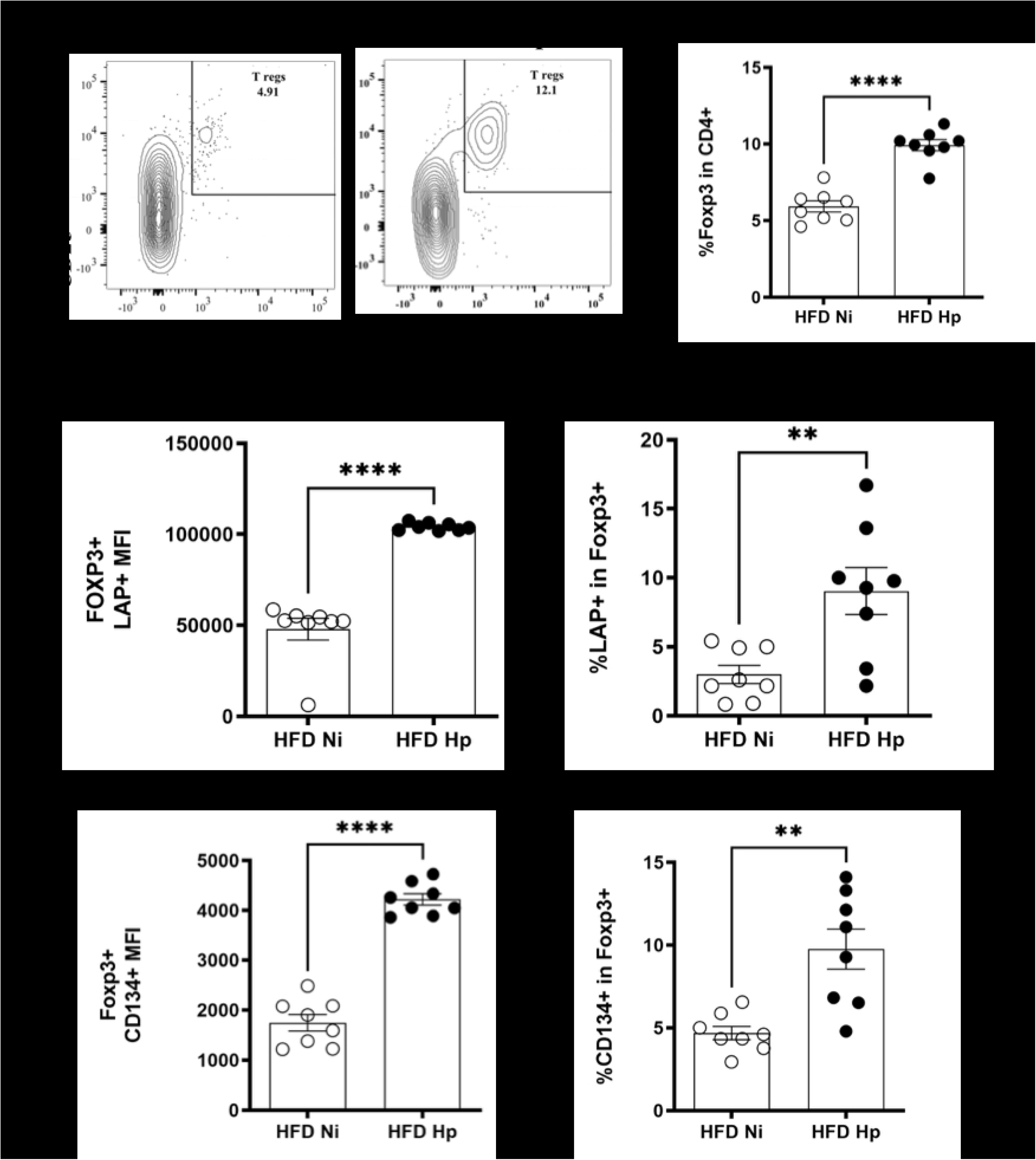
*H. polygyrus* infection induced alterations in number and phenotype of Tregs from EWAT. Representative Contour Plots and percentage of Tregs (A) in HFD Ni and HFD Hp groups, analyzed by flow cytometry. Tregs gated on CD3^+^CD4^+^CD25^+^Foxp3^+^. Mean Fluorescence Intensity (MFI) (B) and percentage (C) of LAP in Foxp3^+^ cells. MFI (D) and percentage (E) of CD134 in Foxp3^+^ cells. n=10 pooled from two experiments performed. Two-tailed t test. ** p<0.01 ****p<0.0001 between HFD groups.

Since the infection altered the expression of functional markers of Tregs, we verified if it could also impact the Treg dysfunction associated with obesity. Adipose tissue Tregs were isolated and used in a proliferation inhibition assay with splenic non-Tregs T cells from LFD-treated animals. Animals from HFD Ni group showed an important dysfunction in Tregs since they were unable to inhibit T cell proliferation at any concentration tested (Fig 4). On the other hand, *H. polygyrus* infection was able to revert this diet-associated dysfunction. Tregs from adipose tissue of infected animals were able to inhibit T cell proliferation at the proportions of 4:1 and 2:1. Together our data showed that *H. polygyrus* infection can modulate different aspects of Treg cells resident in adipose tissue.

**Fig 4.**
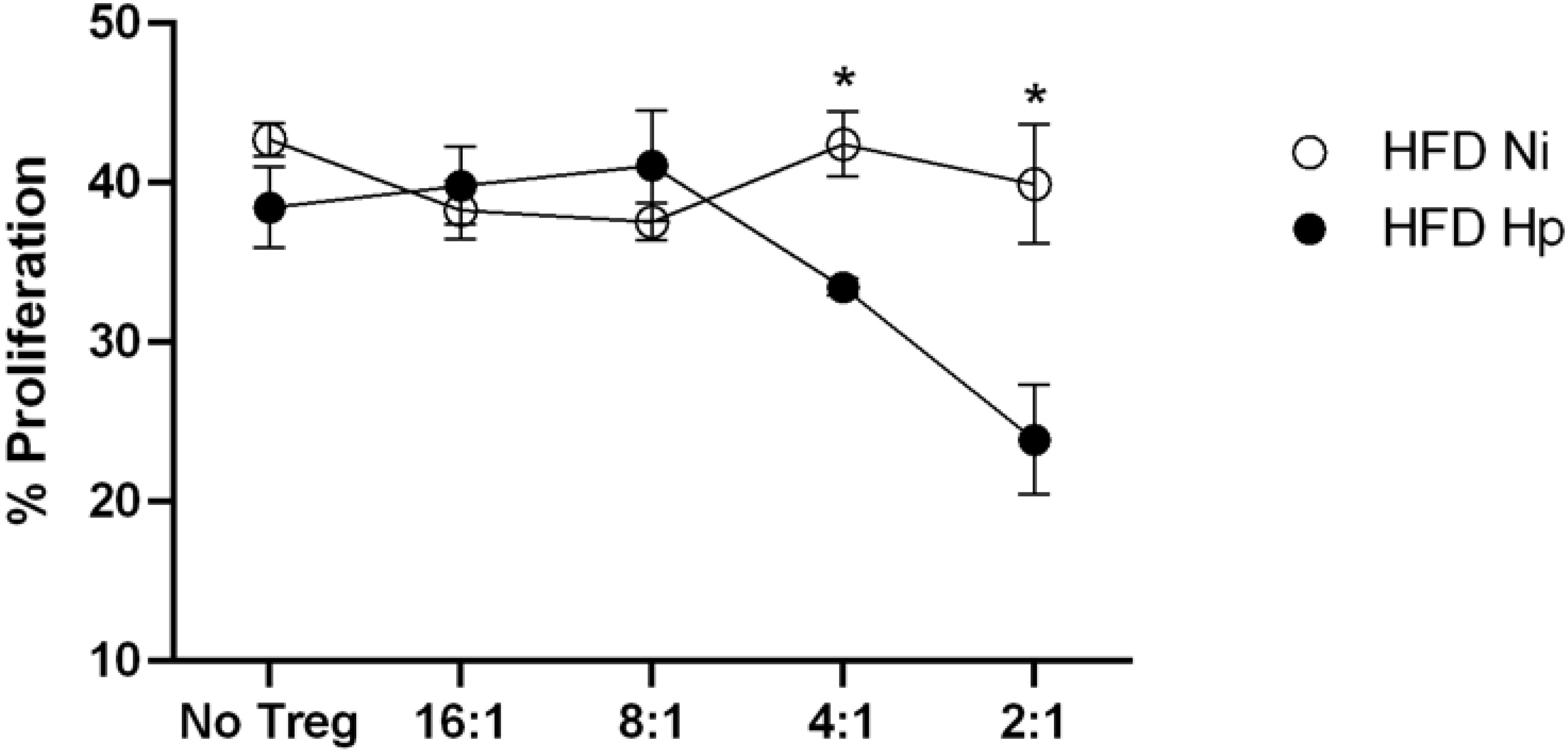
*H. polygyrus* infection prevented loss of function by adipose tissue Tregs. Proliferation percentage, measured by CFSE decay, of non-Tregs cells after co-culture with different concentrations of Tregs. n=10 pooled from two experiments performed. Two-tailed t test. * p<0,05 between HFD groups.

## Discussion

Studies on the effects of helminth infections in metabolic diseases are still emerging and controversial, especially regarding the immunological mechanisms at play in the improvement of metabolic parameters in the host. Our data corroborate previous publications that helminth infections are able to prevent HFD-induced exacerbated weight gain [35, 39, 40], dysfunctional glucose and lipid metabolism [37, 41], and decrease obesity-associated inflammation [35, 36, 38, 40]. Previous reports have associated these effects with a variety of mechanisms such as increased adipose tissue content of eosinophils [37, 40, 60], ILC2 infiltration [37], M2 polarization [35, 36, 40], and PPARg activation [35, 61] indicating that helminth infection might influence metabolic syndrome inducing type 2 immune response polarization. Interestingly, classic inflammation, associated with high levels of IL6 [25], IFNγ [25], TNF [62], IL1ß [63] and CCL2 [64], has been described as necessary for weight gain and development of metabolic syndrome [27, 65], and is believed to be counter regulated by helminth-induced type 2 polarization. Indeed, we also observed higher levels of eosinophils and Th2 cells in the adipose tissue of HFD-treated and infected animals. In addition, we observed decreased frequencies of Th1 and Th17 cells in adipose tissue, which is consistent with the mechanisms speculated to be at play in helminth-associated improvement of obesity parameters i.e., decrease in weight gain and amelioration of metabolic syndrome. Interestingly, SVF cells isolated from HFD Hp animals secreted similar levels of IL17, TNF and IL6 when compared to HFD Ni animals, which might be related to differences associated with ex vivo and in vitro experimentation, or to additional mechanisms beyond inflammation regulation. Of note, we have found that helminth infection was associated with increased levels of adiponectin, IL10 and Tregs in adipose tissue, which may be associated with a plethora of inflammatory and metabolic mechanisms.

Adiponectin is an adipokine that increases insulin function [12, 54, 55] and therefore is very important for glucose metabolism, acting mostly by increasing insulin sensitivity in the muscle [55] and liver [55], and impairing monocyte migration to adipose tissue [55]. Adiponectin is abundant in plasma of healthy individuals, and its levels are negatively correlated with waist circumference, visceral fat weight, triglycerides levels, fast glucose and insulin levels, and also with the development of type 2 diabetes [66 – 68]. The serum concentration of HDL also seems to be positively related to the level of adiponectin [67]. Considering the influence of adiponectin on these parameters, we speculate that lower weight gain, decreased glycemic level and triglycerides, and increased HDL cholesterol observed in Hp infected animals might be associated with increased production of adiponectin. An indirect effect of adiponectin on glucose metabolism in the context of infection was also observed in humans since anti-helminthic treatment was linked to decreased concentrations of this hormone and increased development of insulin resistance [69].

Another homeostatic and regulatory cytokine which was also up regulated by Hp infection was IL10. Although recent studies have suggested that IL10 have a deleterious effect on insulin pathways and weigh gain during experimental obesity [70, 71], when it comes to the context of helminth infection and high fat diet, increased secretion of IL10 is known to control the development of type 2 diabetes [38], improve triglycerides levels [36], and increase sensitivity to insulin [41]. Perhaps, the association between IL10 and type 2 immunity observed in helminth infections, the modified Th2, may shift the impact of IL10 on obesity. In addition, IL10 has also been shown to prevent TNF-induced fat accumulation in the liver [72]. Indeed, we could observe improvement in triglycerides levels, insulin sensitivity, and fat accumulation in the liver in HFD Hp animals that displayed higher production of IL10.

The development of obesity has been linked to a shift from regulatory to pro-inflammatory environment in adipose tissue [27]. Few studies have found that obesity is associated with reduced number [28] and dysfunction of Tregs [57]. Surprisingly, most studies on helminth and obesity have focused on the role of type 2 immune responses, while little is known about Treg cells in the helminth-obesity interaction. We observed that helminth infection was able to modulate the phenotype, and to improve frequency and function of Tregs in the adipose tissue of mice fed HFD. Due to its nature and function, we speculate that Tregs can be directly associated with the regulation of inflammatory parameters like the higher secretion of IL10, and metabolic improvements [28, 73]. Interestingly, the expression of LAP, one of the receptors found to be increased by helminth infection, in Tregs cells is already described to improve glycemic levels, decrease the secretion of inflammatory mediators, impair accumulation of liver fat and reduce hyperplasia in ß-pancreatic cells [47, 74]. CD134, another receptor upregulated by the infection in Tregs, is known to reduce Th1 and Th17 cells differentiation and to sustain Tregs suppression function [75 – 77]. Taken together, these data suggest that Tregs may be implicated in the mechanisms induced by helminth infection in regulation of obesity different than those associated with type 2 polarization.

Taken together, our data show the beneficial effects of helminth infection in early stages of HFD-induced obesity and its associated metabolic dysfunctions. These effects can be attributed to several interrelated or independent events resulting from *H. polygyrus* infection: e.g., increased secretion of IL10 and adiponectin, increased eosinophils frequency, promotion of Th2 differentiation, increased frequency of Tregs, increased expression of Tregs markers like LAP and CD134 and maintenance of Tregs functionality.

Understanding the influence of helminth infection on regulatory mechanisms that may alleviate metabolic syndrome may bring novel approaches to treat or prevent obesity.

## Acknowledgements

The authors thank Dr. Jeffrey Bethony for critical reading of this manuscript and Dr. Luciana Ventura for the support in adiponectin measurement. The authors are also grateful for the technical support from Flow Cytometry platforms in Fiocruz MG and Faculdade de Farmácia da UFMG.

## Supporting information

**S1 Fig. Gate strategy analysis of eosinophils.** Dot/contour plots are representative of the analysis strategy used. Initially gates for leucocytes (SSC-A x FSC-A), single cells (FSC-H x FSC-A) and time (SSC-A x Time) were delimited. Then after mixing the gates using the tool Boolean Gate – Make and Gate, the eosinophils population was determined by CD11b^int^Siglec F^+^.

**S2 Fig. Gate strategy analysis for T lymphocytes subtypes.** Dot/contour plots are representative of the analysis strategy used. After selecting the lymphocytes population (SSC-A x FSC-A), the gate of single cells (FSC-H x FSC-A) was delimited considering only the cells included in the previous gate. If necessary, considering the population from single cells, the gate of time was made (SSC-A x *Time*) to exclude interruptions during acquisition. CD3^+^ cells flowed by CD4^+^ were identified by being T helper cells. This last population was analyzed considering SSC-A x Tbet/Gata3/RORgT, resulting in Th1, Th2 and Th17 populations, respectively.

**S3 Fig. Gate strategy analysis of Tregs.** Dot/contour plots are representative of the analysis strategy used. First the gate of lymphocytes (SSC-A x FSC-A) was delimited, and from it the gate for single cells (FSC-H x FSC-A) was made. From the resulted population CD3^+^ followed by CD4^+^, and CD25^+^ x Foxp3^+^ cells were selected resulting in Tregs. Considering Tregs the gates SSC-A x GITR, CD152, LAP and CD134 were made resulting in the positive population for each marker.

